# Enhanced performance of gene expression predictive models with protein-mediated spatial chromatin interactions

**DOI:** 10.1101/2023.04.06.535849

**Authors:** Mateusz Chiliński, Jakub Lipiński, Abhishek Agarwal, Yijun Ruan, Dariusz Plewczynski

**Author notes:** contributed equally to this work (Mateusz Chiliński, Jakub Lipiński).

## Abstract

There have been multiple attempts to predict the expression of the genes based on the sequence, epigenetics, and various other factors. To improve those predictions, we have decided to investigate adding protein-specific 3D interactions that play a major role in the compensation of the chromatin structure in the cell nucleus. To achieve this, we have used the architecture of one of the state-of-the-art algorithms, ExPecto (J. Zhou et al., 2018), and investigated the changes in the model metrics upon adding the spatially relevant data. We have used ChIA-PET interactions that are mediated by cohesin (24 cell lines), CTCF (4 cell lines), and RNAPOL2 (4 cell lines). As the output of the study, we have developed the Spatial Gene Expression (SpEx) algorithm that shows statistically significant improvements in most cell lines.

## Introduction

The advances in the field of Machine Learning have revolutionized other fields as well. With the increasing computational power and decreasing costs, the predictive power of modern-day deep learning networks allows scientists to apply those methods to various tasks that would be impossible to solve otherwise. Those advances did not omit the genomics field as well. The first attempts to predict the expression solely on the DNA sequence have started just after The Human Genome Project (Lander et al. 2001) - however, they had a vast number of limitations (Beer and Tavazoie 2004; Yuan et al. 2007), and have mostly concentrated on the classical modelling approaches. However, with the expansion of deep learning models, those limitations started to disappear. One of the first major studies on the usage of CNNs (Fukushima 1980) and XGBoost (Chen and Guestrin 2016) started a new era in the prediction of the expression with the introduction of ExPecto (Zhou et al. 2018). Then it continued with the use of CNNs through multiple models, including Basenji2 (Kelley et al. 2018), and finally with the use of transformer-based models like Enformer (Avsec et al. 2021). However, in our study we have decided to take standard approach that is available with the use of CNNs, and expand it further with the change of the input to include also spatial genomic infromation. The ExPecto model that we decided to advance takes 20kbp surrounding the TSS of a given gene and uses expression from that to train a deep neural network to predict the epigenetic factors. Using those factors, the tissue-specific expression is calculated, with a high spearman correlation score. In our study, we have investigated if the epigenetics marks alone are sufficient for the complex task of prediction of the expression - and have given a hypothesis that while they are greatly informative, there is still a place for improvement. We decided that we would like to investigate the effects of the spatial chromatin architecture inside cell nuclei on the expression by investigating the models created with 3D information available and without it. To do that, we have modified the ExPecto algorithm accordingly, so it uses not only the 20kbp region around the TSS but also regions that are linearly distal - but are, in fact, spatially close, thanks to the spatial interactions that are mediated by specific proteins of interest. The overview of the algorithm proposed by us, SpEx (Spatial Gene Expression), is shown in **Figure 1**.

**Figure 1.**
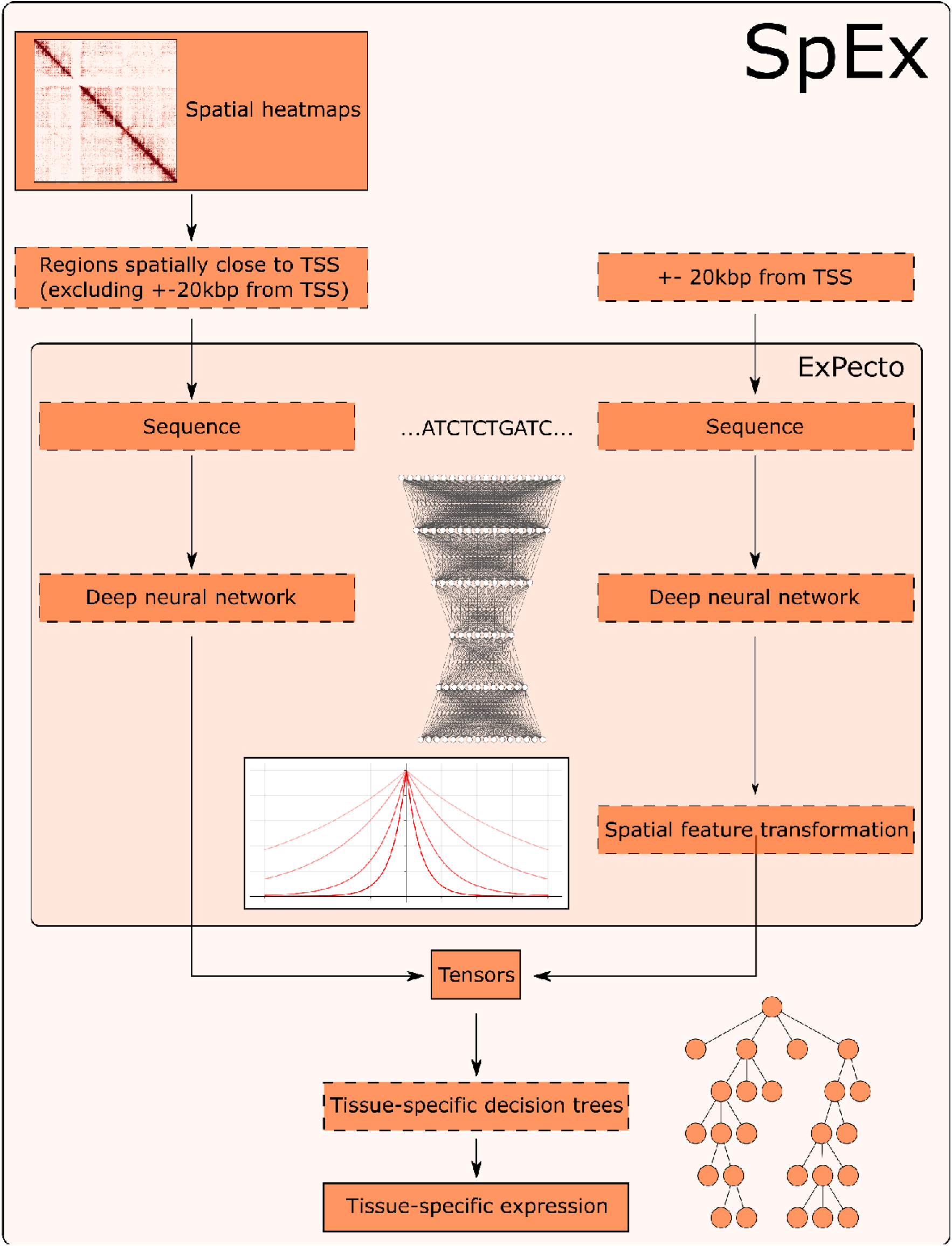
Architecture of SpEx.

To prove the validity of the model, we decided to create an empirical study on how specific protein-mediated interactions are helping in prediction of the gene expression. To do that, we have selected the three most important proteins for loop creation - cohesin, CTCF, and RNAPOL2. The effects of those proteins being unable to bind or be created properly were shown in multiple studies and were the inspiration for asking the question of whether the machine learning models, provided we add 3D information (from interactions mediated by those proteins), will improve.

### Proteins of interest

Cohesin is a protein complex discovered in 1997 (Guacci et al. 1997; Michaelis et al. 1997) by two separate groups of scientists. The complex is made out of SMC1, SMC3, RAD21, and SCC3. However, in human cell lines, SCC3 (present in yeast) is replaced by its paralogues - SA1 (Carramolino et al. 1997), SA2 (Tóth et al. 1999), and SA3 (Pezzi et al. 2000). However, SA3 appears only in cohesin during mitosis (Garcia-Cruz et al. 2010), and we will concentrate in this review mostly on SA1 and SA2 since they are forming cohesin in somatic cells. The complex is extremely important in the proper functioning of the cell nucleus - as is fundamental for the loop extrusion (Davidson et al. 2019), it stabilises the topologically associating domains (cohesin-SA1) (Kojic et al. 2018), allows interactions between enhancers and promoters (cohesin-SA2) (Kojic et al. 2018). The depletion of cohesin in a nucleus removes all the domains (Rao et al. 2017), and completely destroys the spatial organisation of the chromatin. Mutations of cohesin have negative effects on the expression of the genes - e.g. in Cornelia de Lange syndrome (Takahashi et al. 2004; Deardorff et al. 2012) and cancer (Rocquain et al. 2010), where the altered complex is incapable of sustaining its proper function, leading to diseases.

CTCF (CCCTC-binding factor) is 11-zinc finger protein. Its primary function is the organisation of the 3D landscape of the genome (Phillips and Corces 2009). This regulation includes: creating topologically associated domains (TADs) (Guo et al. 2015; Phillips-Cremins et al. 2013; Fudenberg et al. 2016), loop extrusion (Hansen 2020), and alternative splicing (Alharbi et al. 2021). The protein very often works with the previously mentioned cohesin complex, allowing loop formation. CTCF, as a regulator of the genome, binds to specific binding motifs and regulates around that loci. That is why, in case of mutations in the motifs, it might bind improperly, thus allowing disease development. However, not only mutations in the binding sites are disease prone. Mutations in the CTCF protein itself have proven to have a major influence on development of multiple diseases. Some of the examples of diseases inducted by a mutation in the CTCF proteins include MSI-positive endometrial cancers (Zighelboim et al. 2014), breast cancers (Aulmann et al. 2003; Zhou et al. 2004), head and neck cancer (Bornstein et al. 2016).

In eukaryotic organisms, there are 3 common RNA Polymerase complex proteins - I, II, and III (Roeder and Rutter 1969). In this study, we will focus mostly on RNAPOL2, as that is responsible for the transcription of the DNA into messenger RNA (Sims et al. 2004; Orphanides and Reinberg 2002), thus having the biggest impact on the expression of the genes. The mechanisms responsible for creating the RNAPOL2 loops are complex and require not only RNAPOL2 protein but also a number of other transcription factors (Orphanides et al. 1996; Conaway and Conaway 1997). The mutations in those transcription factors have been shown to be linked to various diseases (Aso et al. 1996), including acute myeloid leukaemia (Thirman et al. 1994; Mitani et al. 1995; Rabbitts 1994), Von Hippel– Lindau disease (Whaley et al. 1994; Duan et al. 1995), sporadic cerebellar hemangioblastomas (Kanno et al. 1994), benign mesenchymal tumours (Schoenmakers et al. 1995), xeroderma pigmentosum, Cockayne syndrome, trichothiodystrophy (Scriver 1995), and Rubenstein-Taybi syndrome (Petrij et al. 1995).

### Protein-mediated Interactions

There have been multiple studies showing the spatial landscape created by cohesin-mediated chromatin loops. The first major cohesin ChIA-PET study from 2014 (Dowen et al. 2014) showed the internal organisation of chromatin in the chromosomes. For example, the study provided a list of enhancer-promoter interactions, which can be a starting point for gene expression study.

The next study from 2020 (Grubert et al. 2020) extended the 2014 study and showed that among 24 human cell types, 72% of those loops are the same; however, the remaining 28% are correlated to the gene expression in different cell lines. Those loops are mostly connecting enhancers to the promoters, thus regulating the gene expression. Another interesting insight from this study is that those different profiles of interactions are effective in clustering the cell types depending on the tissue they were taken from.

CTCF, as mentioned above, is responsible for loop extrusion. That is why it is very popular to investigate CTCF-mediated interactions. Once again, like with the cohesin complexes, ChIA-PET is used for obtaining the interactions mediated by CTCF. One of the major studies from 2015 (Tang et al. 2015) shows the genomic landscape among 4 cell lines. They discovered that SNPs occurring in the motif of the CTCF-binding site can alter the existence of the loop - and by that, contribute towards the disease development. They assessed the SNPs residing in the core CTCF motifs and found 70 of those SNPs. Of those, 32 were available from the previously done GWAS studies, and 8 were strongly associated (via linkage disequilibrium) with disease development.

Another study from 2019 (Liu et al. 2019) analysed mutations using 1962 WGS data with 21 different cancer types. Such an analysis, enhanced with the usage of CTCF ChIA-PET data, showed that disruptions of the insulators (that are creating the domains) by motif mutations and improper binding of CTCF (and by that, diminish of the loop) lead to cancer development. Using a computational approach, they have found 21 insulators that are potentially cancerous.

The transcription chromatin interactions, such as the ones mediated by RNAPOL2, are of great interest as well - they control the transcription directly, after all. The study from 2012 (Zhang et al. 2012) showed the RNAPOL2-mediated ChIA-PET interactions on 5 different cell lines, to show the transcriptional genomic landscape. Another study from 2020 (Ramanand et al. 8 2020) performed the same experiments on RWPE-1, LNCaP, VCaP, and DU145 cancer cell lines. They have shown, similarly to in the 2012 study, the spatial interactions based on RNAPOL2, but this time in cancer cell lines. Furthermore, they showed that cohesin and CTCF interactions provide a stable structural framework for the RNAPOL2 interactions to regulate the expression, thus making all of the proteins that we describe in this section crucial for the proper expression of the genes.

Those findings were the main motivation for our analysis - as based on the evidence, the cohesin, CTCF, and RNAPOL2 interactions should give us more information on the genetic expression, thus improving the metrics for the machine learning models. In this work, we present an extension of the ExPecto (Zhou et al. 2018) deep learning model that is enriched with spatial information, thus, as expected, improving the statistical metrics.

## Results

To study those changes, we have gathered the aforementioned 24 cell lines for the cohesin ChIA-PET and 4 cell lines for CTCF and RNAPOL2 binding factors (Dekker et al. 2017; Reiff et al. 2022). They were all mapped to the closest tissue from the GTEx database, where the expressions were taken. The training of the model was performed 1000 times to ensure the statistical significance of the findings. The results for each experiment can be seen in **Supplementary Figure 1**. The highest improvements in the spearman correlation score can be seen in the models that use heatmaps from RNAPOL2 ChIA-PETs. In that case, the metric’s improvement was up to even 0.042 (in RNAPOL2 ChIA-PET GM12878), and the average improvement was 0.016. In the case of CTCF, the highest improvement was also in GM12878, with an improvement of 0.025, with the average improvement over the CTCF study of 0.009. In the case of the cohesin ChIA-PETs, the highest improvement was seen in the K562 cell line, as it totalled 0.020, with an average increase of the correlation score of 0.004. Furthermore, all of the tests were found to be statistically significant, with all the p-values < 10e-11, with exception of two tests: cohesin ChIA-PET KU19, which obtained a p-value of 0.000103, and cohesin ChIA-PET H1, which obtained p-value of 0.01014. The average improvement over the whole dataset was established at 0.0058, and all the grouped sets (cohesin, CTCF, RNAPOL2) were statistically significant at p-value < 10e-31. The cumulative results can be seen in **Figure 2**.

**Figure 2.**
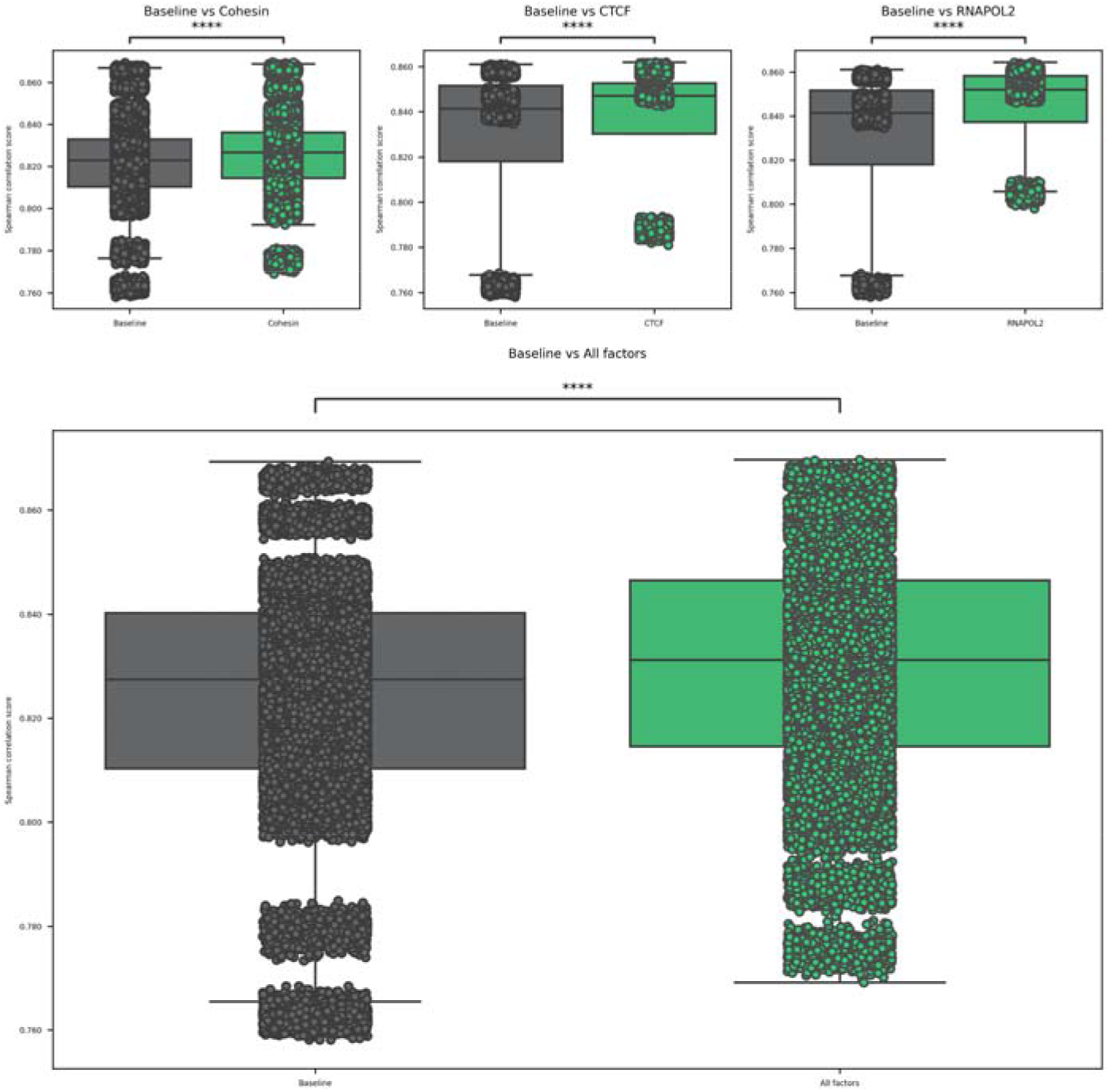
Statistical analysis of the spearman correlation score between the baselines and the experiments grouped by the factor of interest (cohesin, CTCF, RNAPOL2).

Further, to investigate the model in more detail, we compared the residuals of the baseline model with the ones obtained from SpExfor all the proteins. The value of residuals is defined as the difference between observed and predicted values of data, therefore, addressing the quality of the model. We calculated the residuals in the testing set of 990 genes from chr8 for all the models. For the practical analysis, we plotted the density of genes with their associated residual value, which follows gaussian distribution, satisfying the assumption of the normality of the residuals (**Figure 3**). The data is also cross-checked using statistical tests (such as the IFCC-recommended Anderson-Darling test) to ensure it fits a Gaussian distribution. The residual distribution shows the highest improvement in the RNA-POL II compared to the CTCF and Cohesin (**Figure 3. i**).

**Figure 3.**
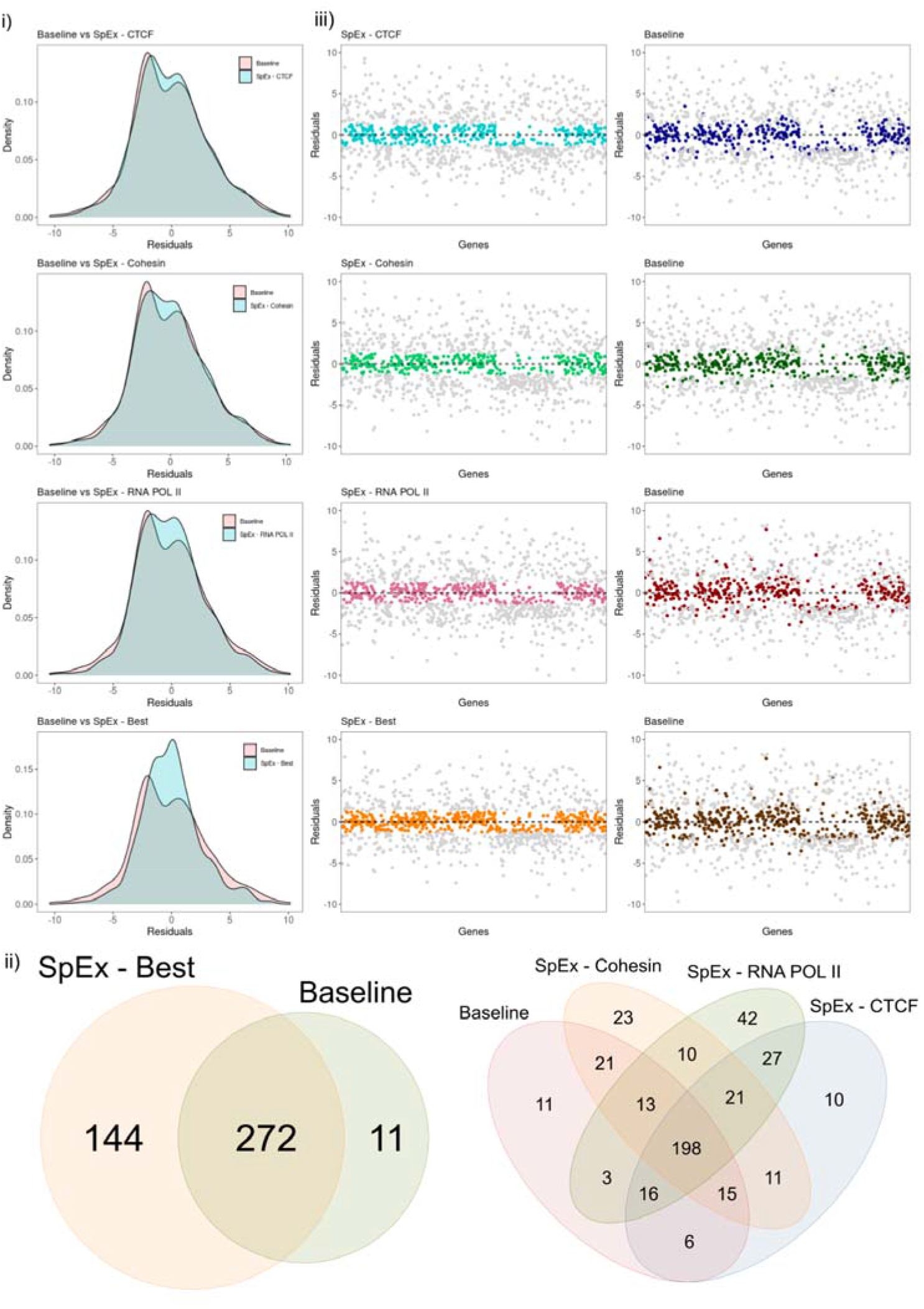
(i) Distribution of residuals for all the protein factors and SpEx-Best, along with the comparison of residual pf baseline. (ii) Venn Diagram of the genes (within cutoff) that are improved by a specific model (iii) Scatter plot of genes (n = 990) with respect to their residual value; highlighted genes are within cutoff (0.5 SD of SpEx), same genes mapped on the baseline.

The architectural proteins - CTCF, Cohesin and RNA POL II, play a diverse role in contributing to gene expression either alone or working together to instruct gene accessibility and expression (Valton et al. 2022; Nora et al. 2012). Therefore, considering that fact, we focused on the residual value of a gene closest to zero by comparing all three proteins named “SpEx-Best”. There is a high density of points close to the origin and a low density of points away from the origin for SpEx-Best compared with the baseline model, which signifies that the gene expression is majorly controlled by the three-dimensional genome structures (**Figure 3. i**).

To investigate the impact of 3D information on gene expression, we conducted a statistical analysis to determine the mean and standard deviation (SD) of the SpEx Best values. We then used this analysis to identify genes that showed the most significant improvement in their expression levels as a result of incorporating 3D information. Specifically, we considered genes within 0.5 SD of the SpEx-Best distribution, which corresponded to a cutoff range of -1.2667 to +1.2667 (**Supplementary Figure 2**). We utilized this cutoff to evaluate the efficacy of our model and found that out of 990 genes, 427 were within this range. Among these genes, 272 were improved in both models, 144 were specific to SpEx, and only 11 were specific to the baseline model (**Figure 3. ii**). Our results emphasize the regulatory role of 3D information in gene expression, which is not captured in the baseline model.

Moreover, we assessed the individual impact of each protein on gene expression and observed that their contributions varied. In particular, RNA POL II showed the highest number of improved genes and thus significantly impacted model performance (**Figure 3. ii**). To further demonstrate the differences, we plotted the value of residuals for each gene for all protein factors and SpEx-Best, highlighting only those genes that fall within the cutoff. We also mapped these highlighted genes (i.e., those within the cutoff of protein factor and SpEx-Best) to the residual of the baseline model (**Figure 3. iii)**. As expected, many genes in the baseline are far from the cutoff and have very high residual values. Therefore we conclude that the proposed model has better efficiency in prediction expression over the baseline model.

To investigate the improvement of the model, we decided to take an example loop that is significant in all three datasets -CTCF, Cohesin, and RNA POL II ChIA-PET. The loop was also required to target gene that had improved prediction score in SpEx over the baseline. The example shows that the gene is spatially close to an enhancer, which plays a key role in altering gene expression. For instance, the enhanced prediction score of the expression of the *TTI2* gene in all three protein factors is due to the fact that the *TTI2* gene interacts with subsequent enhancers that are 20kb apart but close in 3D orientation (**Figure 4**).

**Figure 4.**
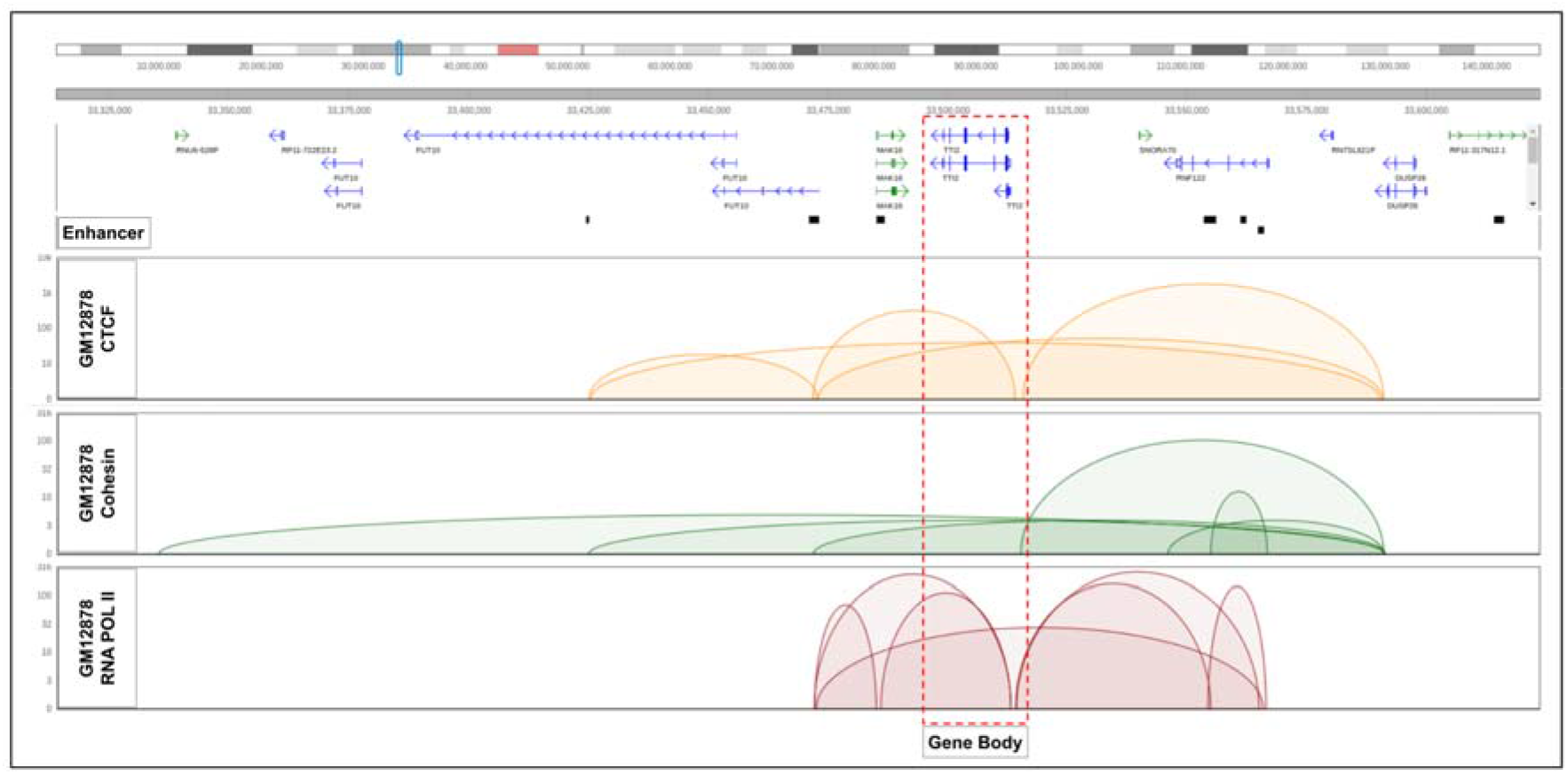
Visualization of the gene *TTI2*, gene body is interacting with the enhancers that are 20kb apart from the TTS in spatial orientation.

## Discussion

In this study, we have shown that chromatin’s spatial structure significantly influences gene expression. To demonstrate that, we have created an algorithm based on the previous work (ExPecto), and added the processing of the spatial heatmaps created by the ChIA-PET experiments. The experiments were performed using 3 different mediating proteins, thus giving us the maps of the interactions involving those proteins. In all 3 cases, the algorithm improved the baseline model, providing us with up to a 0.042 increase in the spearman correlation score. The whole study created 32 experiments, out of which, in 27, we could see improvements. Those findings contribute to the rapid-changing field of three-dimensional genomics, showing that the interactions are indeed required for the proper prediction of the expression - linearly available data, even if we take as many epigenetics factors as in the base ExPecto model (2002), can be still improved with the usage of the spatial data. The next step in the field of prediction of the gene expression is usage of more modern deep learning architectures - e.g. the ones using transformers, like Enformer - and connecting them with the spatial information for the improvement over the baseline models.

## Methods

### SpEx architecture

SpEx, as an extension to ExPecto (Zhou et al. 2018) uses the models described by the authors to generate linear tensors (that are a matrix, where we have 2002 epigenetics features x 10 features showing closeness to the TSS). However, we have added additional spatial information. At the step of generating the final tensors for each of the genes, an additional spatial tensor is added to the linear one. To create it, a number of steps are executed. First, all the contacts that fall out of the linear scope (20 000 base pairs) are taken into consideration. Then, we filter out only the contacts starting or ending near the TSS of the gene, between (TSS, TSS+HiC_resolution), and any other site. Then, only the contacts with a count of at least 0.5 are considered. After getting the regions to predict, that are spatially close to the TSS in an aforementioned way; the ExPecto prediction is run upon those regions. The predicted signal in the regions is summed to ensure that the tensors are uniform in size. That way, we created the tensors that include not only linear information (<20 000 bp) but also consider the signal from the regions spatially close to the TSS of the gene. That way, we are getting a matrix with 2002 epigenetics features x (10 features showing closeness to the TSS + 1 feature representing the regions that are close to TSS in a spatial sense).

The tensors created in that way are saved, as it is computationally expensive to calculate all of them, as both ExPecto and SpEx are calculating them for each of the genes, totalling in 22827 tensors for each of the cell lines. The second step is an actual prediction of the expression. For that, we have used, as in the ExPecto paper, XGBoost (Chen and Guestrin 2016) library. However, we have used different models and parameters. In the case of ExPecto, the model used was GBLinear with reg: linear objective, and we decided to use GBTree with reg:squarederror objective. In the case of SpEx (as the model uses a tree), we have used the tree method of gpu_hist. The full list of parameters used in our model can be found in the code repository.

### Performing the experiments

All the experiments were performed using NVIDIA DGX A100 systems. For each cell line, 22827 tensors were created using one A100 GPU, 8 CPUs, and 128GB physical memory. All the tensors took less than 24 hours to complete with such settings. Following that, each of the cell lines was subjected to the final training 1000 times to ensure statistical significance of the results, meaning that total 53 000 training were completed (32 cell lines + 21 baselines, without spatial information). Individual training operations, in most cases, took up to 5 minutes, and each of the training was assigned one A100 GPU unit, 8 CPUs, and 16GB of physical memory.

### Statistical analysis of the results

From all the experiments was gathered together, and triple statistical testing was performed for each cell line/factor/tissue. We have used welch’s t-test with independent samples with Bonferroni correction from package statannotations (Charlier et al. 2022). The results were also tested for the significance in factor-dependent groups (cohesin, CTCF, RNAPOL2) and all together. The residual analysis was done using an example iteration described in the previous section.

### CTCF & RNAPOL2 datasets

The ChIA-PET CTCF & RNAPOL2 processed data was taken from the 4DNucleome consortium data page (https://data.4dnucleome.org/). The data was obtained there using 4 replicates (2 biological x 2 technical). The pairs were obtained using the ChIA-PIPE (Lee et al. 2020) workflow, which produced pairs for each of the replicates. Then, the pairs were merged and processed using a cooler and juicer to obtain the final .mcool files that were downloaded from the database and used in the SpEx algorithm.

### Processing of cohesin dataset

We gathered the Cohesin ChIA-PET dataset from Encode Portal (https://www.encodeproject.org/) with accession number ENCSR129LGO submitted by Grubert et al. The dataset contains 24 diverse human cell types (Grubert et al. 2020). We merged the replicates and then processed them with the ChIA-PIPE pipeline (Lee et al. 2020) using the default parameters (Linker Sequence = GTTGGATAAG and Peak-calling Algorithm = MACS2). The pipeline generated a high-resolution 2D contact matrix (in .hic file format) along with the annotated chromatin loops with their binding peak overlap. These .hic files were then converted into .mcools files using the hic2cool tool (https://github.com/4dn-dcic/hic2cool) developed by 4DNucleome to obtain the final input for the SpEx algorithm.

## Acknowledgments

This work has been supported by National Science Centre, Poland (2019/35/O/ST6/02484 and 2020/37/B/NZ2/03757); The work has been co-supported by Enhpathy - “Molecular Basis of Human enhanceropathies” funded by the European Union’s Horizon 2020 research and innovation programme under the Marie Sklodowska-Curie grant agreement No 860002 and National Institute of Health USA 4DNucleome grant 1U54DK107967-01 “Nucleome Positioning System for Spatiotemporal Genome Organization and Regulation”. Research was co-funded by the Warsaw University of Technology within the Excellence Initiative: Research University (IDUB) programme. Computations were performed thanks to the Laboratory of Bioinformatics and Computational Genomics, Faculty of Mathematics and Information Science, Warsaw University of Technology, using the Artificial Intelligence HPC platform financed by the Polish Ministry of Science and Higher Education (decision no. 7054/IA/SP/2020 of 2020-08-28).

## Contributions

JL implemented the code of the SpEx algorithm under DP’s supervision. MC updated the algorithm, performed the experiments, and the statistical analysis of the results under DP and JL supervision. All authors prepared the manuscript. AA processed the cohesin datasets and performed residual analysis. MC and JL contributed equally as co-first authors to the whole study. YR provided the CTCF and RNAPOL2 ChIA-PET datasets within the 4DNucleome initiative. DP supervised the whole study. All authors read and approved the final manuscript.

